# Nuclear androgen receptor regulates testes organization and oocyte maturation in zebrafish

**DOI:** 10.1101/159848

**Authors:** Camerron M. Crowder, Christopher S. Lassiter, Daniel A. Gorelick

## Abstract

Androgens act through the nuclear androgen receptor (AR) to regulate gonad differentiation and development. In mice, AR is required for spermatogenesis, testis development and formation of external genitalia in males and oocyte maturation in females. However, the extent to which these phenotypes are conserved in nonmammalian vertebrates is not well understood, because mutations in AR have not been generated in any other species. Here, we generate zebrafish with a mutation in the *ar* gene and examine the role of AR on sexual differentiation and gonad development. We find that zebrafish AR is not required for male sexual differentiation, since a portion of *ar* mutants develop a testis. However, we show that in zebrafish, as in mice, AR is required for the development of secondary sexual characteristics and for proper organization of the testis in males and for oocyte maturation in females. Additionally, we find that zebrafish *ar* mutant males have functional, mature sperm present in their testis, but are infertile due to an inability to release sperm. These findings suggest that AR is required for male sexual development and fertility, but not essential for sexual differentiation in zebrafish. The *ar* mutant we developed will be useful for modeling human endocrine function in zebrafish.

## INTRODUCTION

Androgens are hormones required for proper sex differentiation and development in vertebrates. Androgens act by binding to receptors, such as the androgen receptor (AR), a ligand-dependent transcription factor and member of the nuclear hormone receptor superfamily (1–7). Studies in mice with mutations in the *ar* gene demonstrate that in males, AR is required for spermatogenesis and proper development of the internal reproductive tract and external genitalia (8–10). In females, AR is necessary for folliculogenesis, with impaired oocyte maturation observed in *Ar* mutants mice (11). However, the conservation of these phenotypes outside of mammals is not well understood and has not been explored in zebrafish, a powerful model for the development and function of the human endocrine system.

Zebrafish exhibit unique sex determination and differentiation mechanisms. Laboratory strains lack a sex chromosome and no sex-determining gene has been identified (12,13). Instead, evidence suggests that multiple sex-related genes may be interacting as a network to establish sex, a process described as polygenic sex determination (13). Additionally, all juvenile zebrafish possess a bipotential gonad that appears histologically as an immature ovary (14). In female zebrafish this bipotential gonad matures into a functional ovary, while in males the bipotential gonad regresses and is replaced by a testis (15). Signaling from primordial germ cells (PGCs) has also been shown to be important for female sex differentiation. Ablating PGCs during development produced adult fish that possess a testis but lack sperm (16–19).

Evidence suggests that a growth factor released by PGCs, *bone morphogenic protein 15 (bmp15)*, promotes the female fate by increasing expression of *cyp19a1a*, an aromatase enzyme that converts androgens to estrogens. The subsequent increase in estrogens is thought to promote differentiation of bipotential somatic cells into granulosa cells. Loss of *bmp15* in adults results in female to male sex reversal in zebrafish (20). An additional gene associated with maintaining female fate is *forkhead box L2 (foxl2)*, a transcription factor expressed in ovarian support cells and downregulated in germ cell-depleted zebrafish (17). Other genes associated with the transition from bipotential gonad to testis, including the genes *sry-related HMG box gene 9 (sox9a)* and *antimullerian hormone (amh)* and *doublesex and mab-3 related* (dmrt1) transcription factor (17,21,22). However, whether these genes are upstream or downstream of AR signaling in zebrafish has not been explored.

Expression of zebrafish *ar* during gonad development supports a role for its involvement in sex differentiation. *ar* was expressed in both male and female zebrafish, beginning at 1 day-post fertilization (dpf) and peaking in expression at approximately 25 dpf, during the transition of the bipotential gonad stage into a mature ovary or testis (23,24). Similarly, *ar* was upregulated in testes, compared to the ovaries, at the time of sex differentiation (35 dpf) indicating a potential role of AR in these processes (25).

Furthermore, zebrafish, like many teleost species are vulnerable to androgen exposure during development. Exposure to the antiandrogen fluatamide for 48 hours impaired spermatogenesis (26). Exposure to the androgen receptor antagonist vinclozolin, during juvenile stages, when sex differentiation occurs, produced a population of adults that were mostly female and exhibited reduced fecundity (27). Similarly, rearing zebrafish in the presence of the androgen receptor agonist 17β-trenbolone produced all male adults (28–30). Together, these studies suggest that AR influences sex determination and differentiation in zebrafish.

To investigate the role of AR in zebrafish sexual differentiation and development, we developed *ar* mutant zebrafish using CRISR-Cas technology and compared secondary sex characteristics, gonad morphology, and expression of sex-related genes between wildtype and mutant adults. Our findings suggest that AR is required for proper organization of the testis in males and for oocyte maturation in females, but that AR is not required for sex differentiation of the zebrafish testis.

## METHODS

### Animal care & husbandry

Zebrafish were housed at the University of Alabama at Birmingham (UAB) Zebrafish Research Facility. All procedures were approved by the UAB Institutional Animal Care and Use Committee. Zebrafish embryos were raised in E3B (60× E3B: 17.2 g NaCl, 0.75 g KCl, 2.9g CaCl_2_-H_2_o, 2.39 g MgSO_4_ dissolved in 1 L Milli-Q water; diluted to 1× in 9 L Milli-Q water plus 100 μl 0.02% methylene blue) and housed in an incubator at 28.5°C on a 14 h light, 10 h dark cycle, until 5 dpf. At 5 dpf they were transferred to tanks on a recirculating water system (Aquaneering, Inc., San Diego CA). All zebrafish were from the AB strain (31).

### Generation of *ar* guide RNA and Cas9 mRNA

Guide RNA (gRNA) was generated against the following *ar* target sequence: 5’-GGGTACCACTGCTCGTGCGGAGG-3’. Plasmids pT7-gRNA and pT3TS-nCas9n were obtained from Addgene (numbers 46759, 46757) (32). To generate a vector containing *ar* gRNA, pT7-gRNA was digested simultaneously with BsmBI, BgIII and Sall for one hour at 37°C followed by one hour at 55°C. Oligonucleotides were synthesized by Invitrogen (5’-TAGGGTACCACTGCTCGTGCGG-3’ and 5’-AAACCCGCACGAGCAGTGGTACCC-3’), hybridized to one another in NEBuffer3 (New England Biolabs) and annealed into digested pT7-gRNA using Quick T4 DNA Ligase (New England Biolabs) as previously described (32). The final vector was confirmed by DNA sequencing, linearized with BamHI and used as a template to synthesize gRNA with the MegaShortScript T7 Kit (ThermoFisher). gRNA was purified using the RNA clean & concentrator kit (Zymo Research). RNA concentration was quantified using a Nanodrop spectrophotometer (Nanodrop ND-1000, ThermoFisher). Cas9 mRNA was generated similarly using plasmid pT3TS-nCas9n as template (32).

### Embryo injections

One-cell-stage embryos were injected using glass needles pulled on a Sutter Instruments Fleming/Brown Micropipette Puller, model P-97 and a regulated air-pressure micro-injector (Harvard Apparatus, NY, PL1–90). Each embryo was injected into the yolk with a 1 nl solution of 150 ng/μl of Cas9 mRNA, 50 ng/μl of gRNA and 0.1% phenol red. Injected embryos (F0) were raised to adulthood and crossed to wildtype fish (AB) to generate heterozygous F1 embryos. F1 fish with germline mutations were identified by high resolution melting curve analysis and DNA sequencing to select mutations predicted to cause loss of functional protein. For DNA sequencing, genomic DNA was amplified using primers 5’-GAGTTTGCTGGTCCCATGGA-3’ and 5’-TCCCGTTTGAGAGGTGCAAA-3’, TA-cloned into vector pCR2 per manufacturer’s instructions (ThermoFisher) and then sequenced.

### Genomic DNA isolation

Individual embryos or tail biopsies from individual adults were placed in 100 μL ELB (10 mM Tris pH 8.3, 50 mM KCl, 0.3% Tween 20) with 1 μL proteinase K (800 U/ml, NEB) in 96 well plates, one sample per well. Samples were incubated at 55°C for 2 hours (embryos) or 8 hours (adult tail clips) to extract genomic DNA. To inactivate Proteinase K, plates were incubated at 98°C for 10 minutes and stored at -20°C.

### High resolution melt (HRM) curve analysis

PCR and melting curve analysis was performed as described (33). PCR reactions contained 1 μl of LC Green Plus Melting Dye (BioFire Diagnostics), 1 μl of E× Taq Buffer, 0.8 μl of dNTP Mixture (2.5 mM each), 1 μl of each primer (5 μM; 5’-ATACGGCCGAAGTACTGCTC-3’ and 5’-TACGGATGACGGGTCAGCAT-3’), 0.05 |μl of ExTaq (Takara Bio Inc), 1 μl of genomic DNA, and water up to 10 μl. PCR was performed in a Bio-Rad C1000 Touch thermal cycler, using black/white 96 well plates (Bio-Rad HSP9665). PCR reaction protocol was 98°C for 1 min, then 34 cycles of 98°C for 10 sec, 60°C for 20 sec, and 72°C for 20 sec, followed by 72°C for 1 min. After the final step, the plate was heated to 95°C for 20 sec and then rapidly cooled to 4°C. Melting curves were generated with either a LightScanner HR 96 (Idaho Technology) over a 70–95°C range and analyzed with LightScanner Instrument and Analysis Software (V. 2.0.0.1331, Idaho Technology, Inc, Salt Lake City, UT), or with a Bio-Rad CFX96 Real-Time System over a 70-95°C range and analyzed with Bio-Rad CFX Manager 3.1 software.

### Secondary sexual characteristics & sex ratios

Heterozygous *ar^uab105/+^* adult males and females were bred to each other, offspring were raised to adulthood, genotyped using HRM to examine Mendelian ratios, and secondary sexual characteristics were assayed visually. Zebrafish sex was determined by examining multiple secondary sexual characteristics including body shape, standard length, coloration, presence of genital papilla; characteristics previously shown to be sex specific (34). Fish were anesthetized in 0.2 mg/mL tricaine for approximately 2 min, then measured for standard length with digital calipers, weighed (wet), examined and photographed using a Nikon AZ100 microscope equipped with Nikon DS-Fi1 camera using 1x (anal fin) and 2x (genital papilla) objectives (Figure 1). Due to the feminized secondary sexual characteristics in male ***ar^uab105/uab105^*** homozygotes, sex was confirmed via gonad dissection.

**Figure 1.**
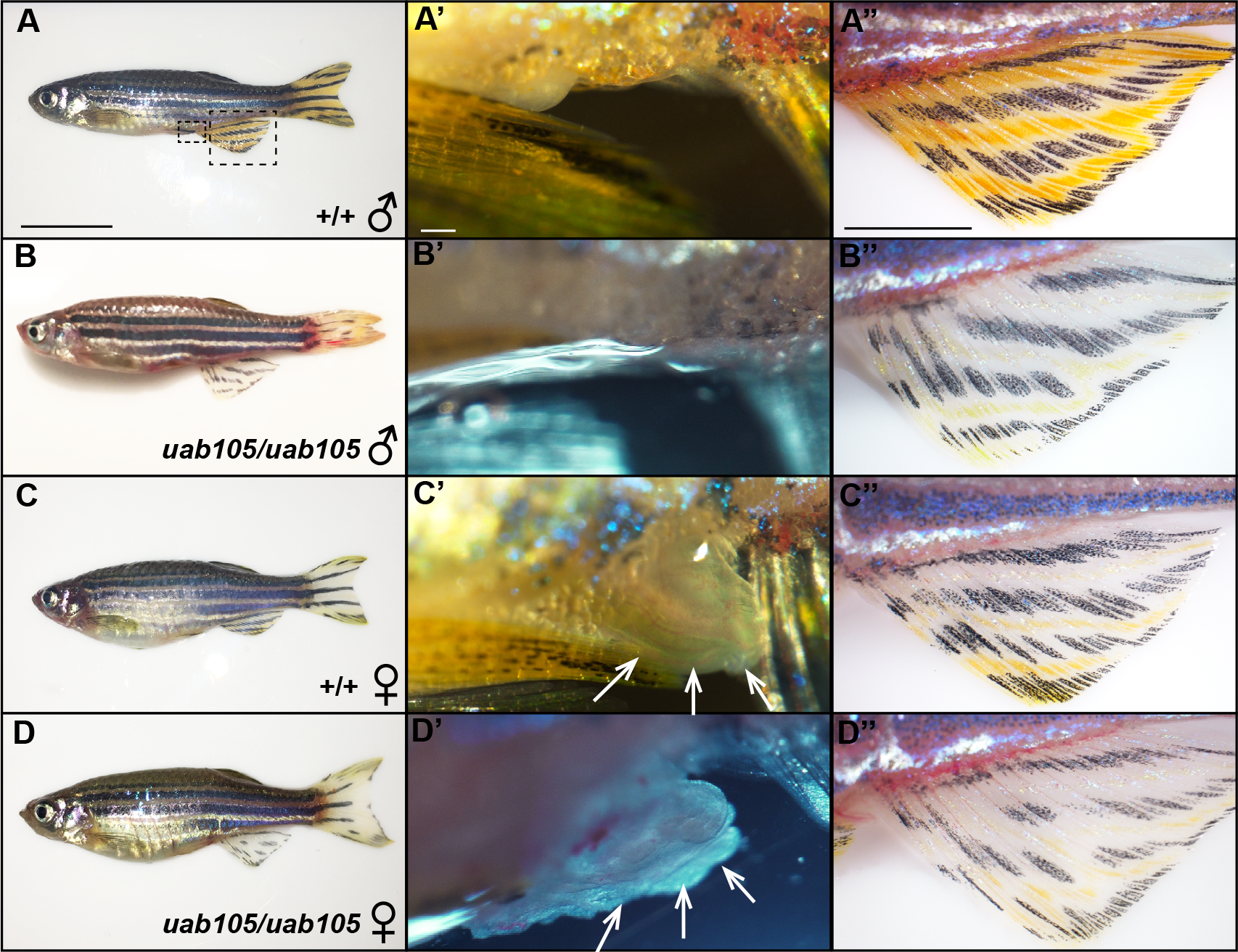
*ar^uab105/uab105^*maleshave feminized body shape and anal fin coloration, whereas *ar^uab105/uab105^* females display normal secondary sexual characteristics. Male wildtype fish have a slender body shape (**A**), lack a genital papilla (**A’**) and have a golden anal fin (**A"**). *ar^uab105/uab105^* mutants have a rounded body shape (**B**), no genital papilla (**B’**), a white anal fin (**B"**), and an internal testis (see Figure 4). Wildtype females (**C**) and *ar^uab105/uab105^* females (**D**) have a rounded body shape with a prominent protruding abdomen, large extended genital papilla (arrows in **C’** and **D’**) and a white anal fin (**C"**, **D"**). (**A’**, **A"**) High magnification images of boxed areas in A, which corresponds to all other similar images. Scale bars: 1 cm (A-D), 100 μm (A’-D’), 1000 μm (A"-D").

### Breeding and *in vitro* fertilization

All wildtype and *ar^uab105/105^* mutant lines were bred on the same schedule, approximately every 2 weeks, to maintain fertility. In order to sex individuals and determine if *ar*^*uab105/105*^ fish were sterile, males and females were set up in breeding tanks with wildtype fish in a 1:2 ratio (mutant to wildtype). In addition, sterility and fecundity was assessed using *in vitro* fertilization (IVF) (31). For IVF, mutant and wildtype fish were anesthetized in 0.2 mg/mL tricaine, patted dry, and their abdomen was gently stroked with blunt-ended forceps to elicit excretion of sperm or eggs. A capillary tube and mouth pipette was used to extract sperm, which upon extraction was activated in 200 μl Hank’s balanced salt solution (HBSS). An additional form of IVF was utilized where male fish were euthanized in iced 0.2 mg/mL tricaine, testes were dissected, mixed with 200 μl HBSS, and minced with a mini mortar and pestle in a centrifuge tube. Expelled egg clutches from a single female were mixed with 50 μl of extracted testis. Sperm obtained from minced testes were visualized and imaged using DIC microscopy on a Zeiss Axio Observer.Z1 microscope with a Zeiss Axio MRc5 camera and 40x objective (Figure 4).

### Histology

Whole-body 9-month old adult wildtype (n = 8, 4 males and 4 females), and ***ar^uab105/105^*** (n = 7, 4 females and 3 males) were selected for histology based on secondary sexual characteristics and IVF results. Fish were anesthetized in iced 0.2 mg/mL tricaine, decapitated and cut just prior to the tail with a razor blade, followed by a 5 day fixation in 4% formaldehyde. Specimens were shipped to HistoWiz Inc. (New York, New York) where they were processed, embedded, sectioned and stained with hematoxylin and eosin (H & E). Processing of tissues into paraffin blocks was accomplished using an automated Peloris II tissue processor (Leica Biosystems) and embedded in Histoplast PE (Thermoscientific). Dehydration steps were as follows: 50% EtOH (15 min, 45°C), 70% EtOH (15 min, 45°C) × 2, 90% EtOH (15 min, 45°C), 90% EtOH (30 min, 45°C), 100% EtOH (45 min, 45°C), 100% xylene (20 min, 45°C) × 2, 100% xylene (45 min, 45°C), Parablock wax (30 min, 65°C) × 2 (Leica Biosystems), Parablock wax (45 min, 65°C). Embedded specimens were cut into 10 μm sections and adhered to charged slides. Slides were heated (65°C, 10 min) in an oven to melt paraffin then rehydrated as follows using a TissueTek Prisma (Sakura): 100% xylene (5 min) × 2, 100% EtOH (2 min) × 2, 95% EtOH (1 min), DI H_2_O (1 min), hematoxlin (1 min), deionized (DI) H_2_O (1 min), defining solution (45 sec) (Define MX-aq, Leica Biosystems), DI H_2_O (1 min), Bluing agent (Fisher Scientific), DI H_2_O (1 min), 95% EtOH (15 sec), eosin (5 sec), 95% EtOH (2 min), 100% EtOH (3 min) × 2, 100% xylene (5 min) × 2.

Bightfield images of sections were obtained using a Zeiss Axio Observer.Z1 microscope with a Zeiss Axio MRc5 camera with 10x (oocytes), 20x (oocytes, testis), and 40x (testis) objectives. Tiled images of oocytes were captured with 10% overlap, and edges were fused using the stitching algorithm of Zeiss ZEN 2 blue edition software. Number of oocytes in stage 1, 2, or 3 in wildtype and ***ar^uab105/105^*** were counted across 9 sections of individual fish (n=3) and averaged (Figure 3, C). Statistical analysis of oocyte stages was completed using a one-way ANOVA with Tukey0027s post-hoc test for multiple comparisons using Graphpad Prism version 7.0 software. A p-value of < 0.05 was considered significant.

### RNA extraction and RT-PCR

Total RNA was extracted from 9-month old adult wildtype and ***ar^uab105/uab105^*** zebrafish following the manufacturer’s guidelines of the TRIzol RNA isolation kit (Life Technologies, CA, US). Dissected gonads were homogenized individually in 500 μL of of TRIzol and total RNA was DNase-treated using a TURBO DNA-Free kit (Ambion, CA, US) to remove genomic DNA contamination. RNA was quantified using a NanoDrop ND-1000 UV-Vis spectrophotometer (Thermo Scientific, MA, US) and quality was determined with gel electrophoresis.

Expression of genes previously shown to be associated with sex differentiation were examined using RT-PCR (Table 1). Genes examined include *cyp19a1a* (24), bone morphogenetic protein 15 *(bmp15)* (35) forkhead box L2 *(foxl2), sry-related HMG box gene 9 (sox9a)*, anti-mullerian protein *(amh)*, and rp113a as a reference gene (17,36). RT-PCR reactions were incubated at 98°C for 1 min, followed by 34 cycles of: 98°C for 10 sec, (60 or 61°C) for 20 sec, and 72°C for 45 sec, then 72°C for 2 min. Products were then run out on a 1% gel and imaged on a Gel Logic 212 Pro (Carestream).

**Table 1.**
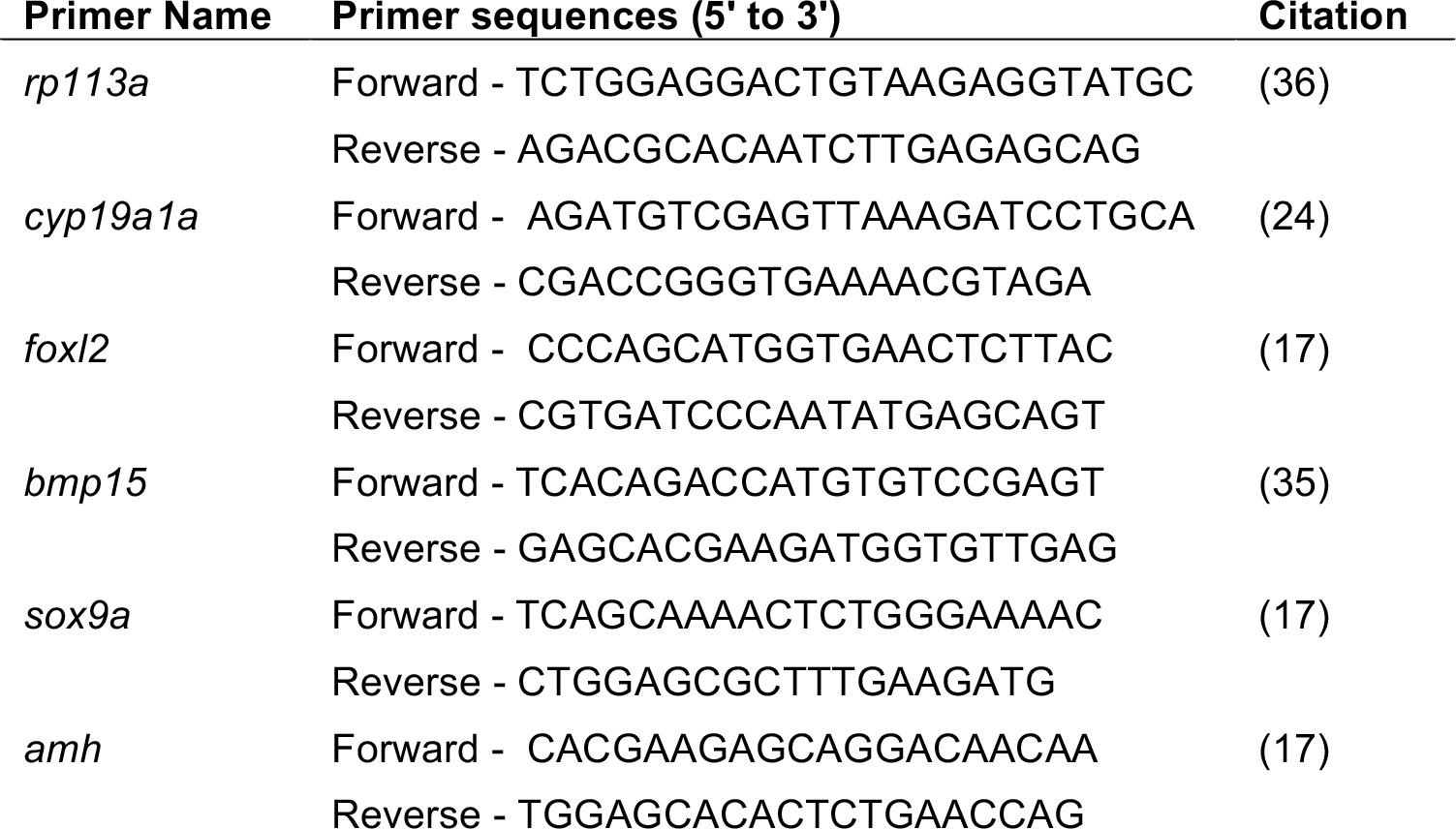
Forward and reverse primer sequences used for RT-PCR.

## RESULTS

### Generation of zebrafish *androgen receptor* mutants

Using CRISPR-Cas technology (32,37), we generated mutations in the zebrafish ***ar*** gene *(nr3c4)* upstream of the DNA binding domain (Figure S1). We identified F1 fish with a 10 base pair (bp) deletion in *ar* leading to a premature stop codon, predicted to have loss of functional protein (Figure S1). Mutants were bred to homozygosity and designated as line *uab105*.

### Male *ar^uab105/105^* have feminized secondary sexual characteristics but lack a genital papilla

To examine the influence of *ar* on zebrafish sexual development, we first assayed secondary sexual characteristics: body shape, coloration, and presence of genital papilla (Figure 1). Females are typically longer than males, have a more rounded abdomen, protruding genital papilla and white colored anal fins. In contrast, males appear smaller than females, have a slender body shape, lack a genital papilla and display a golden anal fin (34).

Heterozygous *ar^uab105/+^* adult fish exhibited wildtype secondary sex characteristics and were easily identifiable as male or female (data not shown). Therefore, to generate homozygous *ar^uab105/uab105^fish*, we mated *ar^uab105/+^* adults. Progeny exhibited the expected Mendelian ratios (1:2:1) of homozygous to heterozygous to wildtype (Table 2). All *ar^uab105/uab105^* had rounded abdomens typical of females and white colored anal fins (n = 67, Figure 1). We observed that a majority of homozygotes had a genital papilla, consistent with wildtype females (n = 53). However, a small portion lacked a genital papilla, consistent with wildtype males, despite exhibiting female-type coloration and rounded abdomens (n = 13, Figure 1B’). Thus, all *ar^uab105/uab105^* adults exhibited female-type rounded abdomen and anal fin coloration, but some lacked a genital papilla.

We hypothesized that the presence of a genital papilla is indicative of an ovary, whereas the lack of a genital papilla correlates to presence of testis. To test this hypothesis, we dissected gonads from *ar^uab105/uab105^* adults. We found a perfect correlation between presence of a genital papilla and presence of ovaries (n = 54 fish), and a perfect correlation between absence of genital papilla and presence of testes (n = 13 fish). Therefore, we indicate males and females based on gonadal sex.

**Table 2.** Mendelian and sex ratios of two populations derived from *uab105/+.* Each population represents the offspring of harem breeding among male and female *uab105/+* heterozygotes. Total number of individual fish within each genotype is indicated, followed by percent of fish with indicated genotype. Numbers of male and females for each genotype are indicated in brackets, with sex based on presence of testis (♂) or ovary (♀).

|  | **+/+** | ***uab105/+*** | ***uab105/uab105*** |
| --- | --- | --- | --- |
| **Population 1** | 58 (25%)<br>[40 ♂, 18♀] | 129 (55%)<br>[84 ♂, 45♀] | 46 (20%)<br>[8 ♂, 38♀] |
| **Population 2** | 24 (25%)<br>[15 ♂, 9♀] | 50 (53%)<br>[29 ♂, 21♀] | 21 (22%)<br>[5 ♂, 16♀] |

We also noticed the ratio of females to males was biased towards females in *ar^uab105/uab105^* compared to heterozygous and wildtype clutch mates (Table 2, Figure 2). In our zebrafish colony, wildtype and heterozygous genotypes exhibited a male bias (58 & 68% males in two separate populations), whereas homozygous *(ar^uab105/uab105^)* exhibited a strong female bias (76 & 83% female, 17 & 24% male) (Table 2, Figure 2).

**Figure 2.**
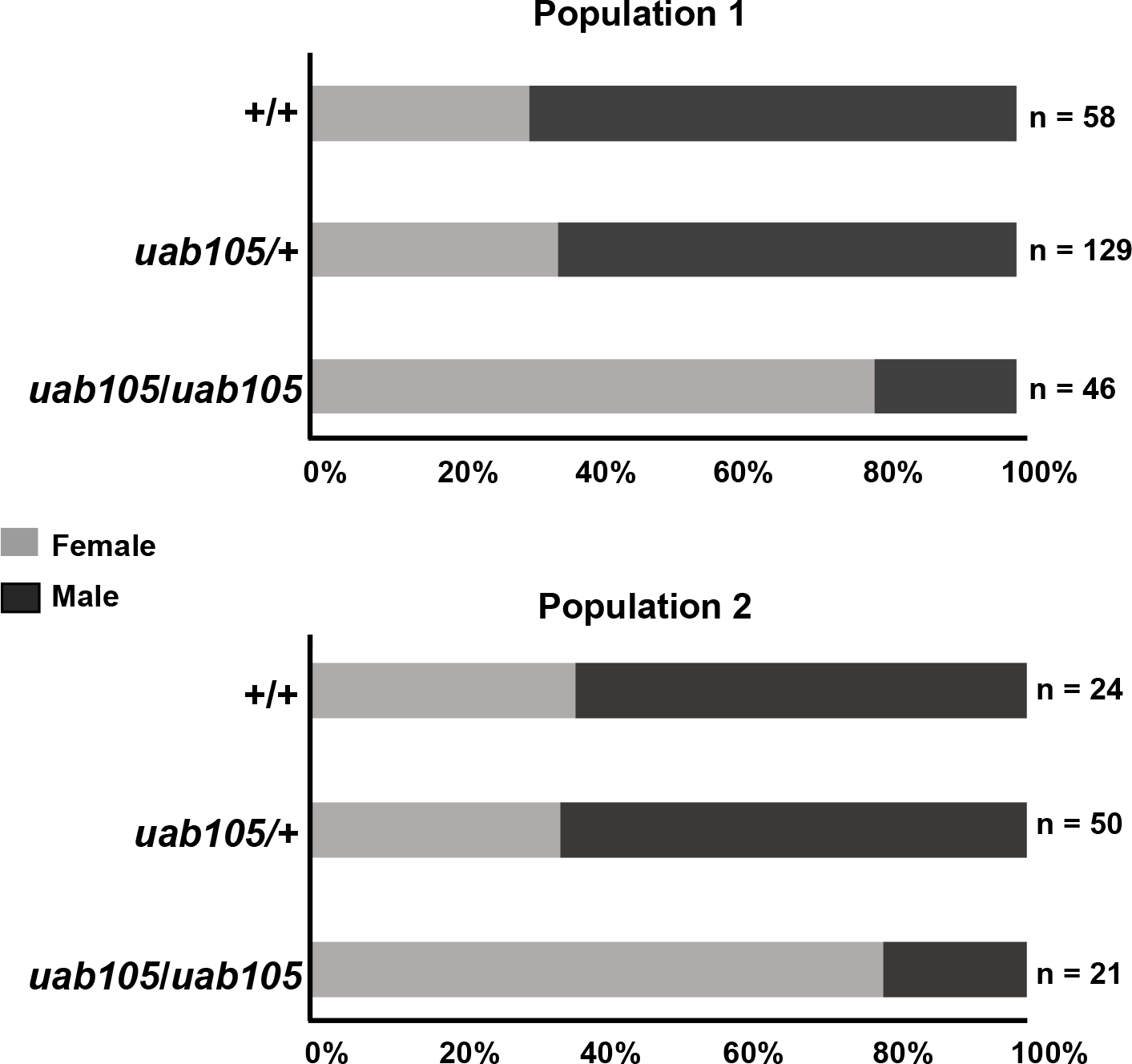
*af^uab105/uab105^* adults are sex biased towards females. Percentages of male and female progeny from crosses between *ar^uab105/+^* males and females. Offspring were observed to have 1:2:1 Mendelian ratios but sex ratios were biased, depending on genotype. See Table 2 for details on Mendelian ratio percentages.

### Male *ar^uab105/105^* produce functional sperm but are unable to ejaculate

To evaluate the influence of *ar* on fertility, we attempted to breed male *ar^uab105/uab105^* (fish lacking genital papilla) with female wildtype under harem spawning conditions (1 male together with 2 females). No homozygous males were successful in breeding with wildtype females under natural spawning conditions (data not shown). This suggests that mutant males either: 1) lack functional sperm, 2) produce functional sperm but cannot release sperm or 3) produce and release functional sperm but fail to induce females to release oocytes. To test whether *ar^uab105/uab105^* produce functional sperm, we attempted to extract sperm from males and fertilize wildtype oocytes *in vitro* using standard, non-surgical and non-invasive procedures (31). After three trials of *in vitro* fertilization, we extracted functional sperm from 100% of wildtype males (n = 3), but failed to extract sperm from any male *ar^uab105/uab105^* fish (n = 13). These results suggest that *ar^uab105/uab105^* males are unable to release sperm.

To determine whether *ar^uab105/uab105^* produce functional sperm despite the inability to release sperm, we dissected testes and immediately visualized them under a microscope, without fixation. We observed sperm with heads and tails in both wildtype and *ar^uab105/uab105^* fish (Figure 3), although mutants had qualitatively fewer numbers of sperm per field of view compared to wildtype.

**Figure 3.**
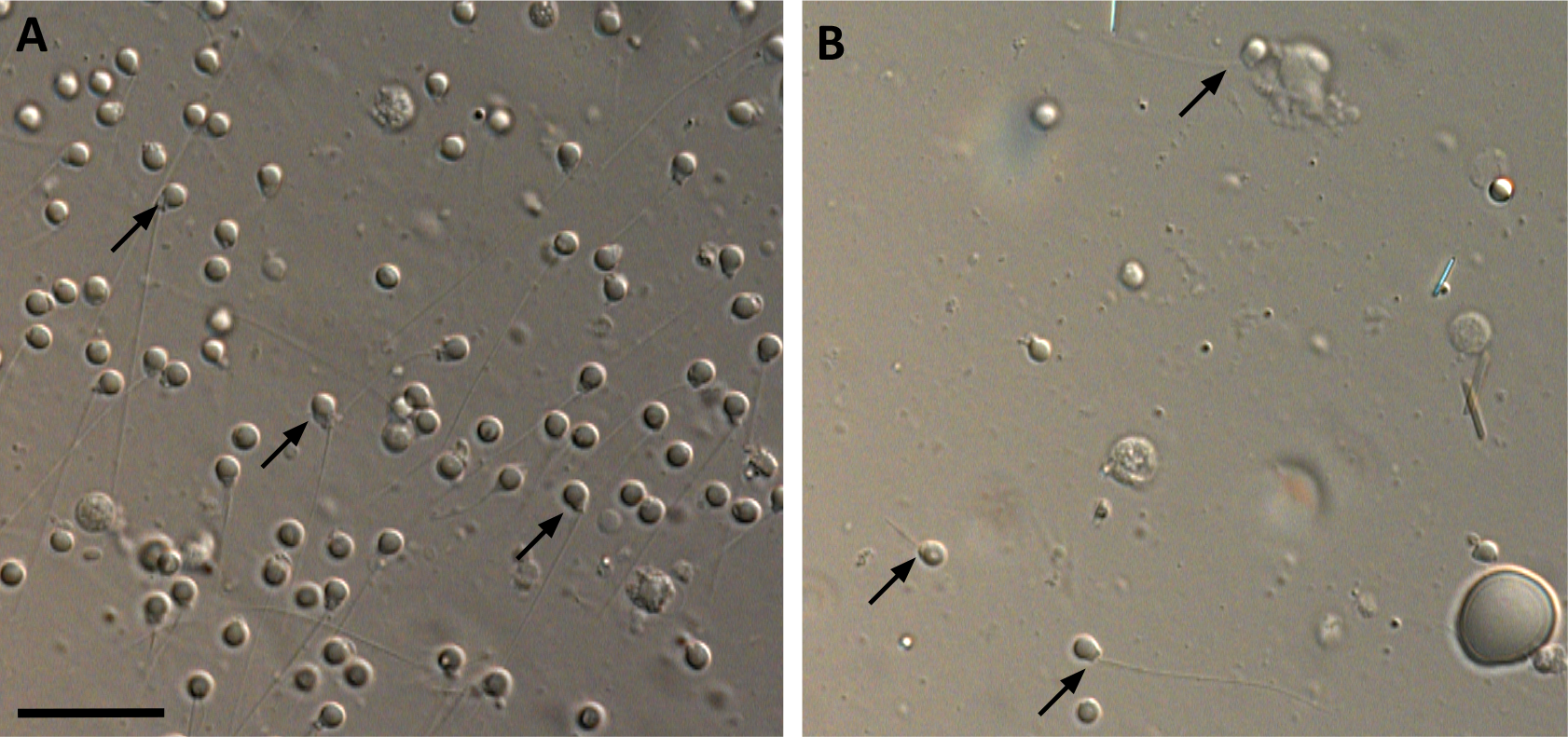
*ar^uab105/uab105^* estis contain mature sperm. Wildtype (**A**) and *ar^uab105/uab105^*(**B**) mature sperm (arrows) from dissected testis. Scale bar = 20 μm

To test whether ar^uab105/uab105^ sperm were able to fertilize oocytes, we mixed *ar^uab105/uab105^* testes extracts with wildtype oocytes and assayed the number of fertilized embryos. We found that sperm from dissected, minced testis from *ar^uab105/uab105^* were capable of fertilizing wildtype oocytes (Table 3). Fertilized oocytes appeared grossly normal through 4 dpf, at which point we euthanized and then genotyped each larva. All larvae were heterozygous *ar^uab105/+^* as expected (n = 24). Therefore, we conclude that *ar^uab105/uab105^* males produce, but are unable to release, functional sperm.

**Table 3.** Sperm from surgically removed testis of *ar^uab105/uab105^* can successfully fertilize wildtype oocytes *in vitro*. Each wdiltype egg clutch was divided and exposed to surgically removed and minced testis from either one wildtype or one mutant fish. Number of 4 dpf larvae derived from wildtype or *uab105/uab105* sperm are indicated for each clutch of oocytes.

|  | Clutch 1 | Clutch 2 | Clutch 3 | Clutch 4 | Clutch 5 | Clutch 6 |
| --- | --- | --- | --- | --- | --- | --- |
| wildtype | 0 | 16 | 0 | 4 | 12 | 3 |
| *uab105/uab105* | 0 | 0 | 2 | 22 | 0 | 0 |

### Male *ar^uab105/uab105^* have disorganized seminiferous tubules and aberrant cyst formation

Because male *ar^uab105/uab105^* were unable to release sperm, we assayed testes for morphological defects. We performed H & E staining on tissue sections from *ar^uab105/uab105^* testes and observed disorganized seminiferous tubules compared to wildtype testes (Figure 4; representative images from n = 3 fish). In wildtype testes, we observed distinct seminiferous tubules, surrounded by a basement membrane, filled with developing sperm organized into readily demarcated cysts with mature sperm (spermatozoa) congregated in the lumen cavity, consistent with previous findings (38). In contrast, *ar^uab105/uab105^* males lacked seminiferous tubule structure and exhibited disorganized cyst formation with no central pool of spermatozoa. Sperm of similar developmental stages (based on diameter of nucleus) were clustered together in cysts, but the cysts were not clustered into tubules as in wildtype testes (Figure 4, compare A” & B”). Additionally, sperm cysts were qualitatively smaller in mutants compared to wildtype. Our results suggest that AR is required for proper testicular organization in zebrafish.

**Figure 4.**
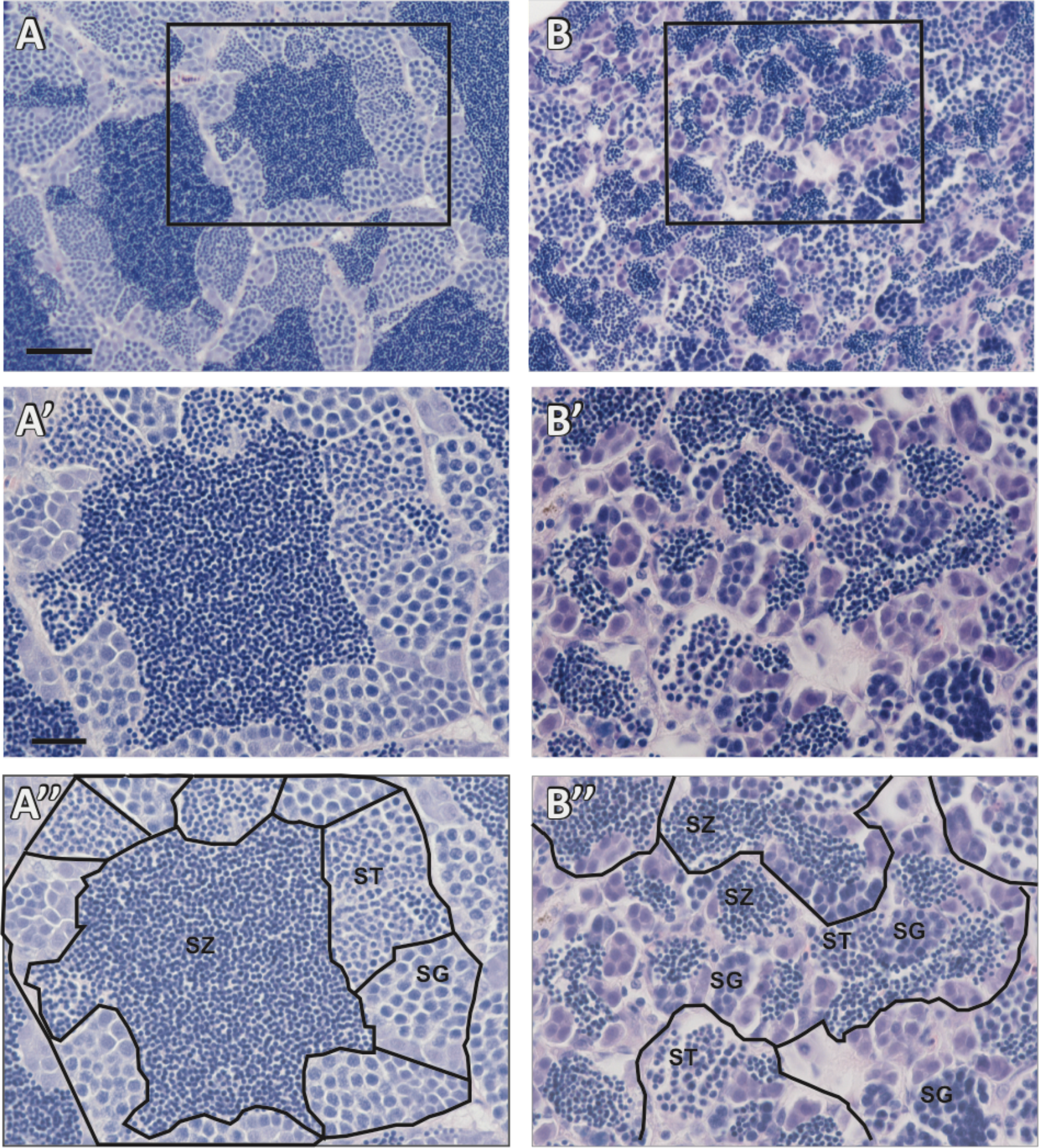
*ar^uab105/uab105^* testes are disorganized and lack seminiferous tubule structure. H & E staining of adult male wildtype (**A**) and *ar^uab105/uab105^* (**B**) sectioned testis. Boxed area in **A** and **B** represent **A’**, **A”**, **B’** and **B”,** respectively. Wildtype testis contain spherical seminiferous tubules surrounded by a basement membrane with cysts of developing sperm surrounding mature sperm in the tubule lumen, whereas *ar^uab105/uab105^* males lack seminiferous tubule organization and cyst structure. **(A”, B”)** Tubules and cysts from A’ and B’ are outlined in black and spermatogenesis stages are indicated: spermatogonia (SG), spermatids (ST), spermatozoa (SZ). Scale bars: 50 μm (A, B), 20 μm (A’, B’, A’’, B’’).

### Female *ar^uab105/105^* have decreased fecundity and fewer mature oocytes

In contrast to *ar^uab105/uab105^* males, *ar^uab105/uab105^* females were fertile and able to breed with wildtype and *ar^uab105/+^* zebrafish. However, fecundity appeared to be reduced in *ar^uab105/uab105^* mutant females compared to wildtype, with only approximately 1 out of 4 natural spawning attempts between wildtype males and *ar^uab105/uab105^* females producing fertilized embryos (data not shown). To determine whether there were morphologic defects in mutant ovaries that might reduce fecundity, we performed H & E staining on tissue sections from wildtype and *ar^uab105/uab105^* ovaries. The overall architecture and organization of ovaries was similar between wildtype and mutant females (Figure 5; representative images from n = 4 fish per genotype). Ovaries from wildtype and mutant fish contained oocytes of different stages from less mature stage I (primary growth stage) through stages II (corticol alveolus stage) to mature (vitellogenic) stage III oocytes. Stages were determined by the appearance of variably sized cortical alveoli, prominence of the vitelline envelope and accumulation of yolk proteins (39). However, the ratio of stage I to stage III oocytes was different, with wildtype containing more stage I oocytes compared to stage III (Figure 5C). Wildtype fish contained on average 21.1 ± 7.4 stage I oocytes and 36.2 ± 11.4 stage III oocytes, compared to 62.8 ± 18.9 stage I oocytes and 7.6 ± 2.5 in mutants (p = 0.0001, ANOVA with Tukey’s test for multiple comparisons, n = 3 fish per genotype, 9 tissue sections analyzed per fish, mean ± standard deviation). There was no significant difference in stage II oocytes between wildtype and *ar^uab105/uab105^* mutants (27.5 ± 6.4 versus 21.1 ± 4.0, p = 0.12). These results indicate that AR influences oocyte maturation in zebrafish.

**Figure 5.**
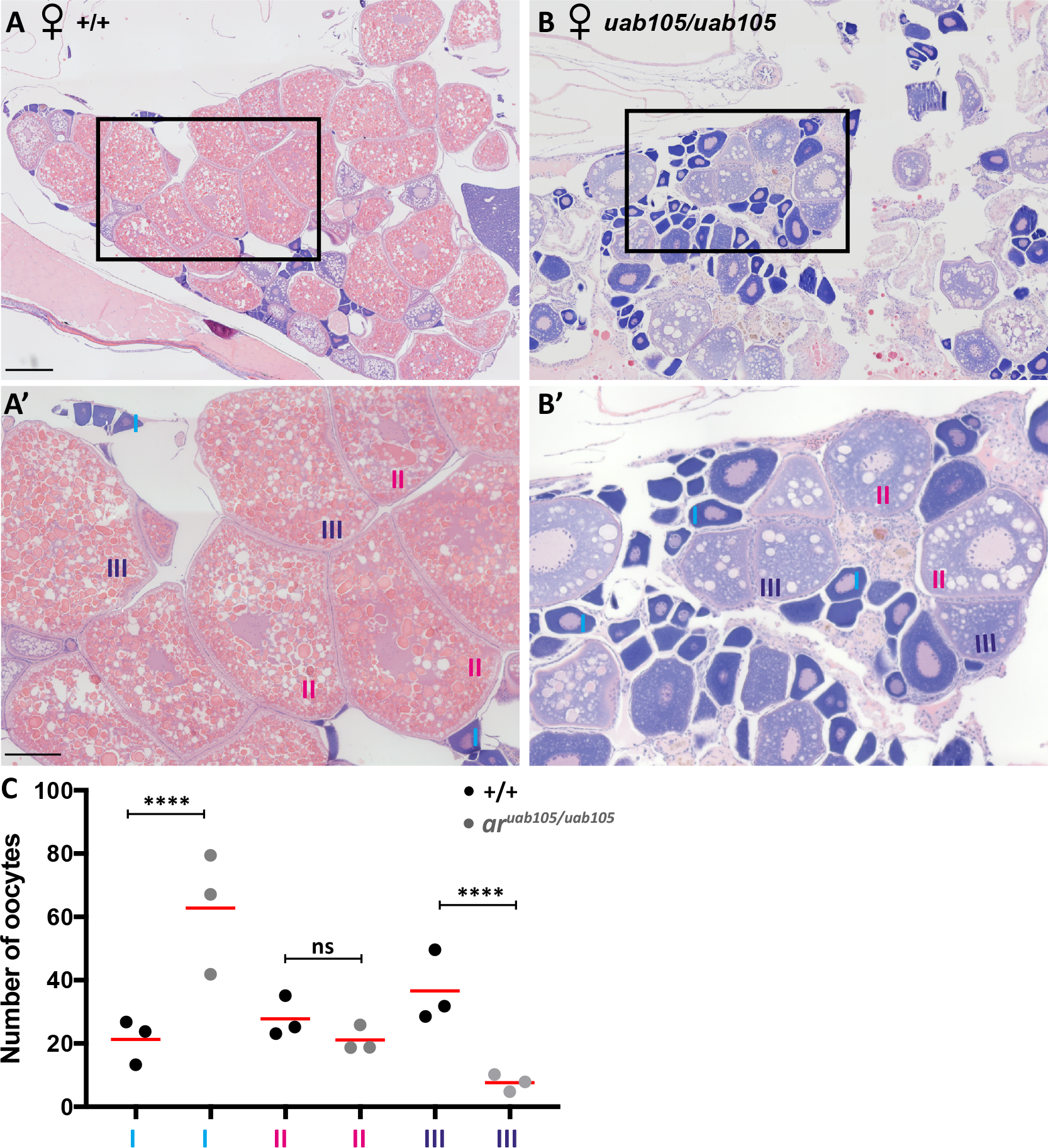
*ar^uab105/uab105^* ovaries have fewermatureoocytes than wildtype. H & E staining ofadult Pemole wild type(**A**) ond *ar^uab105/uab105^* (**B**) ovary sections. (A’, B’) High magnification images of boxed in area in A, B. Wildtype ovaries consist of a majority of mature late-stage III, followed by stage II oocytes, with I oocytes in less abundance. *ar^uab105/uab105^* ovaries contain primarily early-staged I and II, with almost no stage III oocytes. Scale bar: 200 μm (A, B), 50 μm (**A’**, **B’**). (**C**) Quantification of oocyte number according to stage in wildtype and mutants. Each dot represents average number of oocytes per histological section per fish (n= 3 fish per genotype, 9 sections per fish). ****, p < 0.0001 one way ANOVA. ns, not significant.

### Genes associated with sex differentiation are similarly expressed in wildtype and *ar^uab105/1o5^* mutants

To determine whether nuclear androgen receptor acts upstream or downstream of genes known to influence sex differentiation and ovary function in zebrafish, we employed a candidate approach and assayed the expression of five genes known to influence sex differentiation and ovary function in zebrafish. We used RT-PCR to compare gene expression in ovaries and testis dissected from adult wildtype and *ar^uab105/uab105^* mutants (Figure 6, n = 3 fish per sex/genotype) and used the reference gene *rpll3a*, which is expressed in all tissues, as a control. Two genes associated with ovary differentiation and function, *cyp19a1a* and *foxl2* (17,24,40,41), were expressed in ovaries of both wildtype and *ar^uab105/uab105^* mutant females, with faint expression in wildtype and mutant males. Two genes, *bmp15* and *sox9a*, associated with gonad development and maintenance, were expressed in both ovaries and testis of all wildtype and *ar^uab105/105^* mutants. The *amh* gene, expressed by granulosa cells in mature ovaries, was expressed in ovaries from both wildtype and *ar^uab105/uab105^* mutants, with little or no expression in males. We conclude that these five known sex differentiation genes are expressed normally in *ar^uab105/uab105^* mutant gonads. Our results suggest that AR is not required for sex determination or gonad differentiation, but is required for proper gonad function in zebrafish.

**Figure 6.**
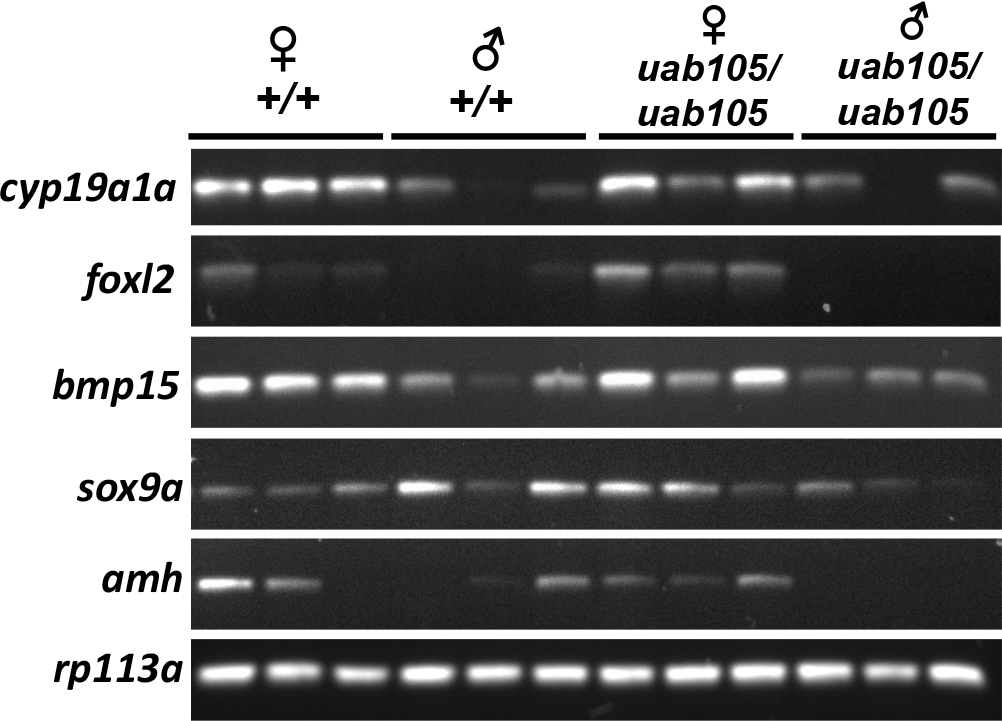
Expression of genes associated with sexeal differentiation in wildtype and *ar^uab105/uab105^* zebrafish. Expression of genes in wildtype (+/+) compared to *ar^uab105/uab105^* mutant females and males assayed by RT-PCR. Expression patterns of all
genes is similarly expressed in wildtype and *ar^uab105/uab105^* mutants testes (♂) and ovaries (♀). ***rp113a*** is included as a reference gene expressed uniformly across samples.

## DISCUSSION

Our data suggests that nuclear AR is not necessary for male sex determination and development in zebrafish, since a portion of *ar^uab105/105^* develop as males with a testis. This was unexpected, considering that exposure to potent androgens or genetic ablation of *cyp19a1a*, an aromatase enzyme responsible for estrogen production, was reported to fully masculinize zebrafish (20,28–30). One possibility is that *ar* is not the only androgen receptor in the zebrafish genome. ZIP9 (*slc39a9*), a member of the zinc transporter subfamily, was shown to bind androgens with high affinity in Atlantic croaker (42). In addition, a calcium and amino acid sensing G-protein coupled receptor (GPRC6A) has been shown to transduce non-genomic effects of androgens in mice (43). Both of these genes are conserved in zebrafish, and therefore could be involved in androgen-dependent testis determination and differentiation. Alternatively, the presence of testes and functional sperm in *ar^uab105/105^* mutant males could be because the *uab105* allele does not result in a complete loss of function of AR, but rather is a hypomorph. Because the targeted mutation in the ***ar*** gene created a frameshift and premature stop codon before the DNA and ligand binding domains (Figure S1), the *uab105* mutant is expected to be a null allele. However, we found that *ar^uab105/105^* populations were biased towards females, consistent with pharmacologic data in other fish species. Female sex biased populations were also reported in juvenile guppies exposed to AR antagonists (44). Together with our results, this suggests that AR influences sex determination, but is not required for male sex determination in zebrafish.

Regardless of whether *uab105* is a hypomorph or true null allele, our results demonstrate that AR is required for the development of some male secondary sex characteristics and for proper testis organization in zebrafish. This is consistent with *Ar* mutant mice, which exhibit female secondary sex characteristics, such as a shorter external genito-anal distance and a decrease in body weight (more similar to females) (8). Additionally, gene expression studies suggest that AR acts downstream of *foxl2*,*bmp15*, *cyp19a1a*, *sox9a* and *amh* and is not required for sex determination or gonad differentiation, but is required for proper gonad function in zebrafish.

Our results indicate the *ar^uab105/105^* were infertile because although they produce functional sperm, they are unable to release sperm. This could be due to improper formation of the duct that transports sperm to the cloaca or due to a blockage within the duct. In mice, AR is important for development of the internal reproductive tract, including the vas deferens and epididymis (8–10). This could indicate a conserved role of AR in the formation of the analogous tubule ducts in zebrafish.

In wildtype zebrafish testes, distinct seminiferous tubules are filled with developing sperm organized into spermatogenic cysts with mature sperm (spermatozoa) centralized in the lumen cavity (38). We found that testes of *ar^uab105/105^* were structurally disorganized, with a lack of seminiferous tubules containing developing sperm surrounding a lumen filled with mature spermatozoa (Figure 4). The mechanisms leading to this disorganization defect are unknown. ***Ar*** mutant mice were reported to have smaller testes that appeared to lack normal organization and were described as thinner and less cellular, compared to wildtype mice. In addition, ***Ar*** mutant mice had impaired spermatogenesis, with developing sperm arrested at the pachytene spermatocyte stage with an absence in spermatids and mature spermatozoa (8,9). Therefore, the presence of mature sperm in ***ar*** mutant zebrafish is in contrast to findings in mice. This could indicate that AR has slightly different roles in spermatogenesis between mammals and zebrafish. One possibility is that in zebrafish, other androgen receptors such as ZIP9 or GPRC6A contribute to androgen-dependent spermatogenesis. Alternatively, *ar^uab105/105^* mutants might be hypomorphic and demonstrate a partial but not complete loss of AR function. Future studies using multiple mutant *ar* alleles will be necessary to clearly evaluate gene function.

In contrast to males, our results suggest that AR is not required for the development of female secondary sexual characteristics or ovaries. However, *ar^uab105/105^* females appeared to have decreased fecundity, with oocytes being released less often during breeding and IVF trials, compared to wildtype females. Histological examination of ovaries in consistently bred females suggests *ar^uab105/105^* mutants have a decreased number of mature, stage III oocytes compared to wildtype (Figure 5). This finding is consistent with results in frogs and mice, where androgens are the primary physiological regulator of oocyte maturation (11,45). *Ar* mutant mice had decreased fecundity and fewer follicle cells, symptoms of premature ovarian failure (10,46). In frogs, treatment with human chorionic gonadotropin, an inducer of ovulation, resulted in high testosterone levels that was attenuated by the AR antagonist flutamide, suggesting that androgens act through AR to induce oocyte maturation (45). High estrogen levels delay oocyte maturation in zebrafish (47). Loss of AR could result in higher levels of circulating androgens, which would be converted to estrogens by aromatase *(cyp19a1a)*, resulting in higher estrogen levels and thus delayed oocyte maturation. While this manuscript was being prepared, an *ar* mutant zebrafish was reported (48). The authors found that homozygous *ar* mutant males exhibited reduced courtship behavior compared to wildtype. However, they did not examine sperm function in males or sex bias, fertility or gonad morphology in males or females.

The zebrafish *ar* mutant developed here is a valuable tool for studying AR function. This mutant will also be important for exploring the molecular and cellular mechanisms by which androgen-like endocrine disrupting compounds influence embryonic development, organ formation and function.

## ACKNOWLEDGEMENTS

We thank R. Swanson and J. L. King for excellent technical assistance, J.M. Parant for help with high resolution melting curve analysis, S. N. Romano, J. P. Souder and H.E. Edwards for support and suggestions, and S. C. Farmer and staff at the UAB Zebrafish Research Facility for animal care. This work was funded by grants from the NIH (K12GM088010 to C.M.C., R01ES026337 to D.A.G.), by a Faculty Research Grant from Roanoke College (to C.S.L.) and by start-up funds from UAB and the Department of Pharmacology & Toxicology.

---

**Figure S1.**
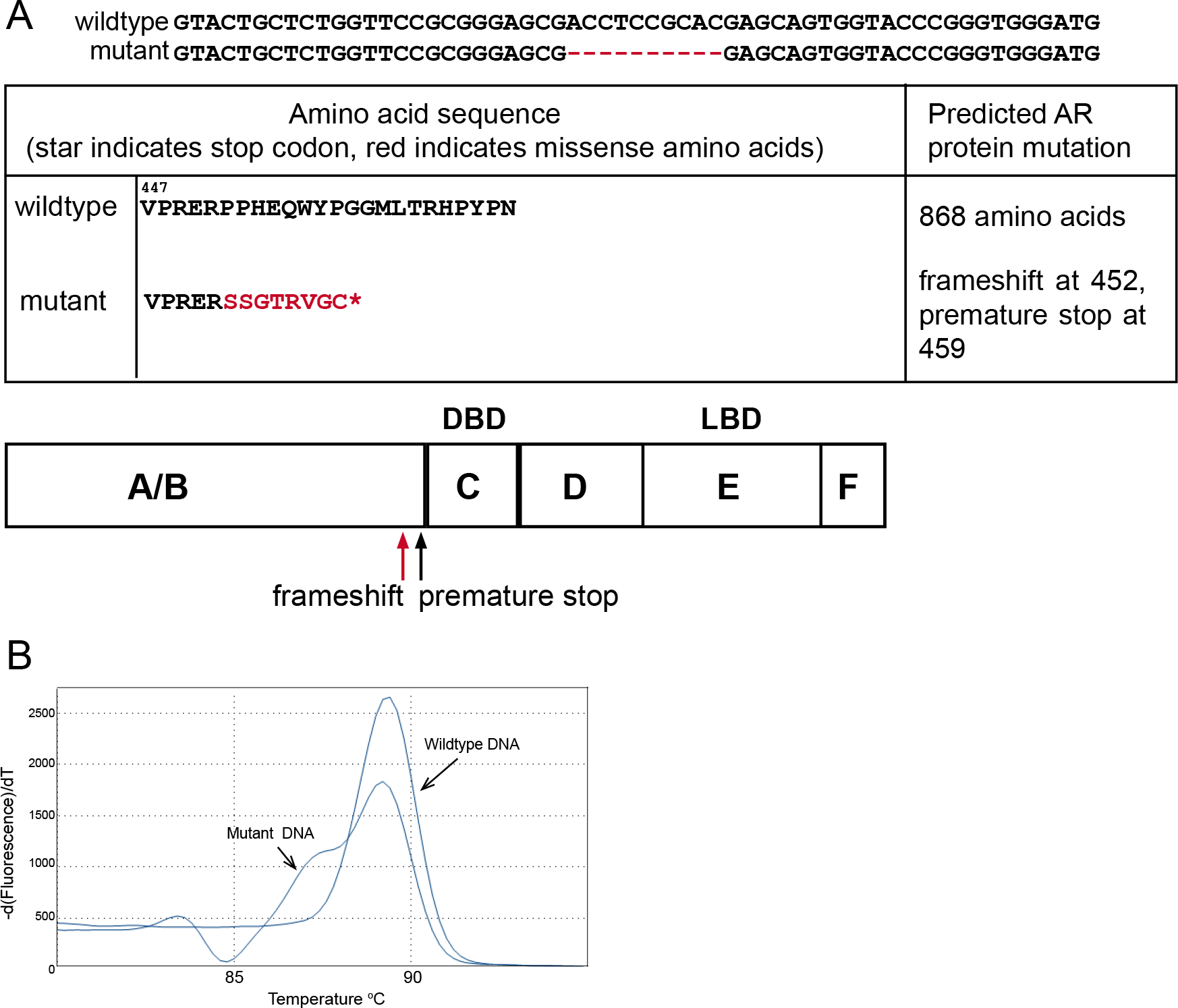
Generation of ar mutant zebrafish. **(A)** Genomic DNA of *ar^uab105^* zebrafish contains a 10 basepair deletion in the ar coding region, resulting in a premature stop codon in the AR protein. Nucleotide deletions are shown as red dashes, amino acid mutations are in red. Map indicates site of frameshift mutation and premature stop codon (DBD, DNA binding domain; LBD, ligand binding domain). **(B)** High resolution melting curve analysis was used to distinguish mutants from wildtype. Curves represents DNA amplified from a wildtype AB or *ar^uab105^* mutant zebrafish (arrows).

